# Optical tomography reconstructing 3D motion and structure of multiple-scattering samples under rotational actuation

**DOI:** 10.1101/2024.11.27.624241

**Authors:** Simon Moser, Mia Kvåle løvmo, Franziska Strasser, Judith Hagenbuchner, Michael J. Ausserlechner, Monika Ritsch-Marte

## Abstract

Optical Diffraction Tomography (ODT) has emerged as a powerful tool for imaging biological cells in a non-invasive and label-free manner. However, conventional approaches using ODT by varying the illumination are plagued by the missing cone problem, which introduces ambiguity and deteriorates the axial resolution in the reconstruction. Although utilizing object rotation has the potential to yield isotropic resolution, experimental control or prior retrieval of the rotational parameters is challenging. In this work, we demonstrate ODT of multiple-scattering samples undergoing variable rotational motion, unlocking the potential for isotropic resolution in non-contact systems. We introduce a comprehensive reconstruction method to jointly retrieve both sample and rotational motion in 3D. An interferometric setup enables the recording of amplitude and phase data while the object is rotated in a non-contact manner around one or more chosen axes in an acoustofluidic device. We evaluate the tomographic reconstruction performance of the method for clusters of micro-beads and highlight its suitability for biomedical application beyond single cells, demonstrating high-resolution reconstruction of dense cancer spheroids containing more than 100 cells.

## 1. Introduction

In this work, we present an approach for performing tomographic imaging of rotating objects in non-contact settings, where the precise rigid motion of the sample is uncertain. We demonstrate how implementing this method in Optical Diffraction Tomography (ODT) [1–3] makes previously challenging samples accessible for label-free 3D tomographic reconstruction. ODT is an imaging method to determine the refractive index (RI) distribution of transparent microscopic objects. In contrast to fluorescence-based methods, ODT relies on the light scattered elastically by the sample structures themselves to generate contrast. Quantitative RI information correlates to the protein distributions in biological cells and, therefore, provides insight into biochemical and morphological properties that are crucial for diagnosis and disease monitoring [4, 5]. As a label-free imaging modality, ODT has proven to be an excellent tool for studying live specimens in a minimally invasive manner. It is, however, usually limited to single cell imaging.

In order to obtain a reasonable 3D reconstruction in ODT, access to views from multiple directions or sample orientations is typically necessary. To acquire the scattered field components from multiple views, a common strategy consists in varying the illumination on a static object [6–19]. This modality maps well to optical microscopes and allows for precise control over the illumination angles. However, modifying the illumination on a static object is limited by the finite numerical aperture (NA) of illumination and detection optics, which leads to the well-known missing-cone problem. To alleviate this issue, incorporation of prior information about the sample structure in the reconstruction process can be pursued by learning-based methods such as classical optimization [10–12, 14, 18, 19], machine learning [20–22] or combinations thereof [23–28]. Although such approaches can reduce the effects of the missing cone problem, certain spatial frequency components remain inaccessible, which unavoidably leads to ambiguity. Instead of scanning the illumination angles on a stationary object, acquiring multiple views by object rotation has potential to yield reconstructions of significantly higher quality as arbitrary object views can be collected. ODT has been performed based on object rotation by mechanical rotation of capillaries [29, 30], and by non-contact methods in fibre-optical traps [31], optical tweezers [32, 33], by flow-based methods [34–36], and in devices based on dielectrophoresis [37] and acoustic forces [38]. Furthermore, combinations of illumination-angle scanning and object rotation have also been demonstrated [33, 39].

Contact-less manipulation in conjunction with optical tomography may offer a fully non-invasive and label-free approach, which is particularly important for studying developing *in vitro* models such as cells, spheroids and organoids, as these models can be sensitive to mechanical influences [40].

The price to pay with non-contact strategies is that the uncertainty in the sample orientations is more pronounced than under mechanical fixation. Standard tomographic reconstruction algorithms require the sample motion to be retrieved prior to the reconstruction. Uncertainty in the sample orientation deteriorates the resolution and quality of the reconstruction and introduces artifacts and distortions. In response to contact-less rotational actuation, the object typically follows a certain trajectory given by experimental conditions, which is difficult to control or predict. A multitude of strategies have been proposed to retrieve the motion of objects rotated in a non-contact manner for ODT. Several approaches rely on prior knowledge of the sample to retrieve the motion, which restricts the strategy to samples with certain shapes or features [31, 35, 41–43]. A common assumption is that the rotation axis is fixed and the angular velocity is constant [38, 41, 43], which may lead to inaccuracies particularly for arbitrarily shaped objects. A few approaches consider corrections to the variable angular velocity [31, 36]. In all of these methods the detection of the angular orientation has limitations and the approaches were restricted to single cells or small specimens.

To enable high-resolution reconstructions of rotated objects, precise motion control or recovery is crucial. How well-defined the motion can be in non-contact approaches varies between modalities, but some degree of rotational variation or translation for objects of arbitrary shape is inevitable. In devices using bulk acoustic waves for sustained or transient object rotations, variation in angular velocity and in rotation axis, in addition to translation is typical [44–46], and recovering the unknown object structure from such data becomes a challenging problem. We recently demonstrated an optimization-based reconstruction algorithm capable of recovering the unknown motion and object parameters of acoustically reoriented objects imaged from multiple angles with OCT [46]. In this case, the motion between 3D OCT data was recovered and the resolution in the reconstructed object reflectance, attenuation and RI maps demonstrated a great accuracy of the algorithm.

In the current work, we propose a comprehensive approach for 3D motion recovery for ODT of arbitrary objects undergoing a rotational motion that is not precisely known. Our approach can deal with multiple-scattering objects and additional experimental unknowns. It employs a gradient-based optimization model that leverages very general prior information on object and motion, such as assuming a rigid body and continuous and periodic motion. The approach is explained schematically in Fig. 1: The object of interest is illuminated from a single direction while being held and rotated in a sustained manner, in our case by acoustic actuation (a). The scattered field components are collected using an optical microscope and recorded interferometrically (i.e. in amplitude and phase). The rotational motion is to a good approximation periodic, but it has some *a priori* unknown irregularities in the trajectory, as it can exhibit non-constant angular velocity and variations of the rotation axis (b). The uncertainty in these parameters make existing approaches for reconstruction difficult or impossible to apply. We utilize prior information in the form of regularization and initialization in our inverse scattering problem, where we enforce the rotational parameters to form a smooth trajectory (c) and solve for the object and motion jointly using gradient-based optimization. We validate our method experimentally on various micro-bead clusters of known properties and apply it then to increasingly challenging biomedical samples, finally demonstrating isotropic high-resolution reconstructions of dense cancer cell clusters containing many cells.

**Fig. 1.**
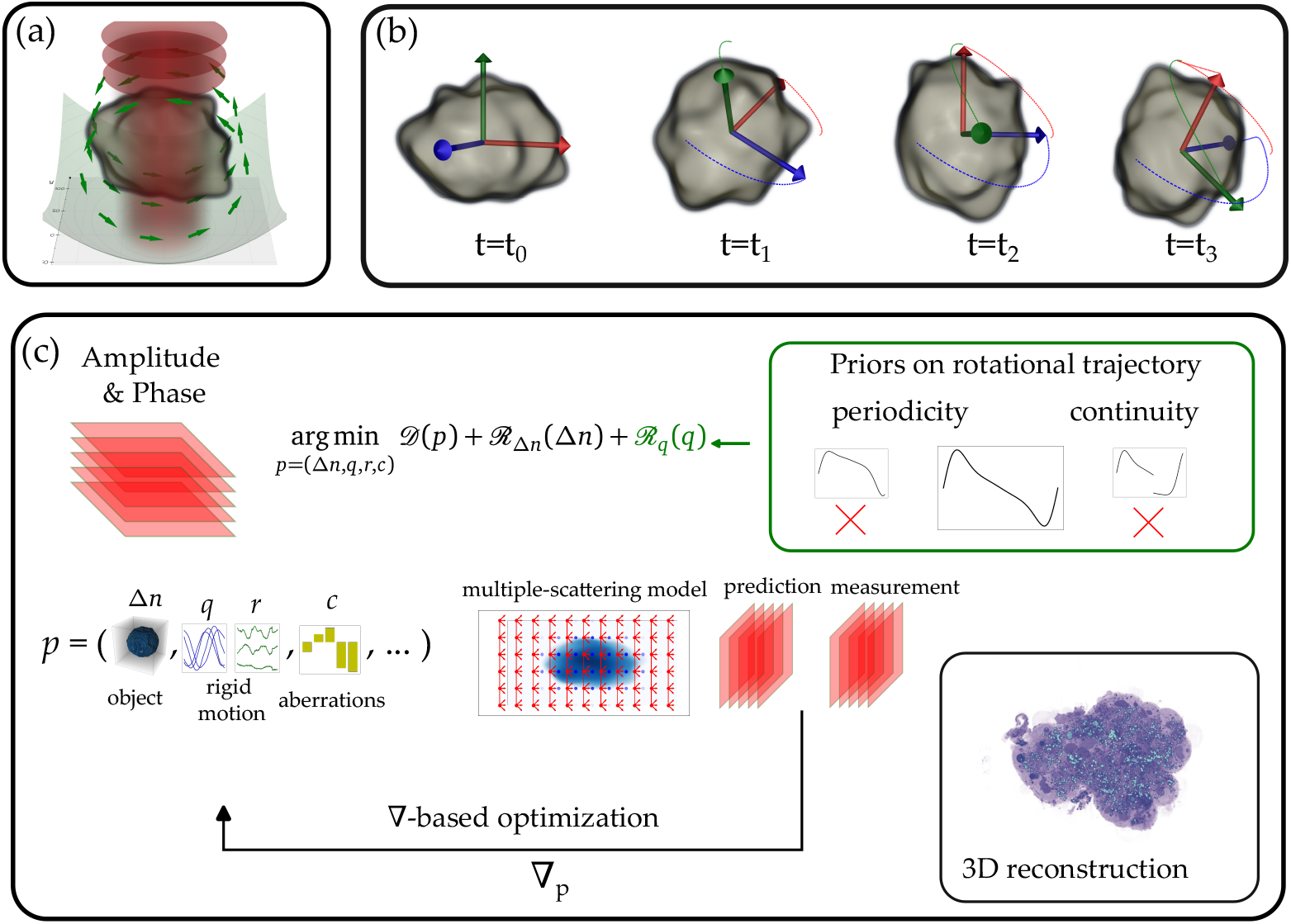
Concept: (a) The sample is trapped and rotated by means of acoustic forces, while scattered field components are recorded. (b) The rotational motion of the sample is not precisely known *a priori*, but exhibits continuity and is approximately periodic. (c) These properties can be incorporated as priors in the inverse scattering problem, which is then solved by means of gradient-based optimization retrieving refractive index, rigid motion and calibration parameters (aberrations).

## 2. Methods

### 2.1. Experimental implementation

A schematic depiction of the experimental setup, comprising optical microscope, acoustofluidic device, and interferometer, is shown in Fig. 2. We trap and rotate the sample using the acoustofluidic chamber developed in [45]. Controllable 3D acoustic trapping is realized by coupling bulk acoustic waves in three orthogonal directions into a microfluidic chip and generating standing waves in the fluid channels. We use lithium niobate (LiNbO_3_) transducers and operate them at the 1st-5th harmonic frequency (3-20 MHz), choosing the trapping wavelength to be comparable to the targeted sample size. The sample is introduced into the center of the chip at the intersection of all acoustic waves and illuminated through the optically transparent vertical transducer. By tuning the frequency, relative amplitude and phase of the transducers we can induce sustained rotation and reorientation of levitated samples. Our acoustofluidic device is also tailored to induce rotations around orthogonal object axes, both perpendicular to the optical axis. For more details we refer to Supplement 1 and [45]. The acoustofluidic device is mounted on a commercial inverted light microscope (Nikon ECLIPSE Ti2-E). We illuminate the sample through the vertical transducer using a collimated beam from a diode laser (TOPTICA iBeam smart *λ* = 639 nm). The scattered field components are collected with a water-immersion objective (Nikon CFI Apochromat LWD Lambda-S 40× NA 1.15) and follow a standard path through the microscope. To obtain phase-stability with trans-illumination through the transparent transducer, a home built common-path interferometer is attached to an output port of the microscope. The interferometer, adapted from [47], generates interferograms which we record with a camera (mvBlueFOX3-2071). To retrieve amplitude and phase, we use off-axis interferometry, which does not require phase stepping.

**Fig. 2.**
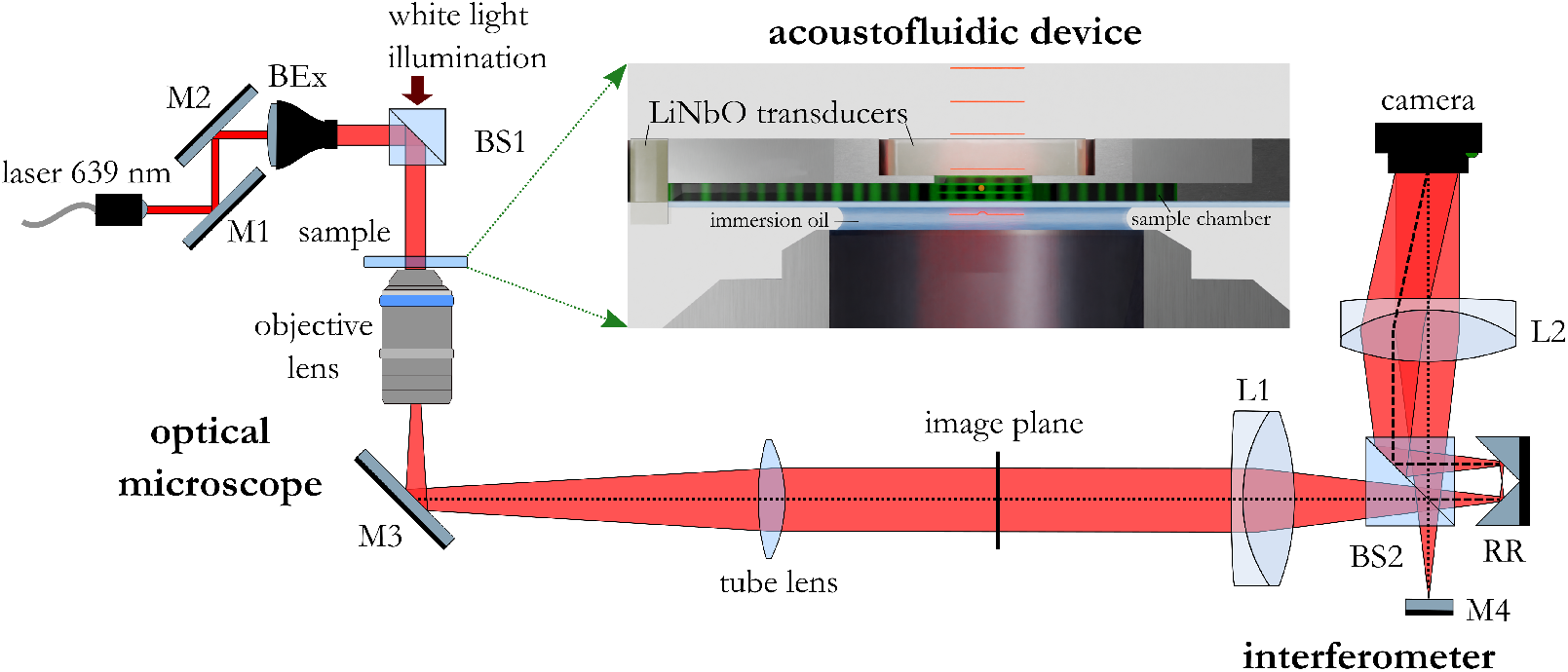
Experimental setup: The sample rotated in the acoustofluidic chip is illuminated by light emitted from a fiber-coupled diode laser (*λ* = 639nm) and the scattered light is collected by a water-immersion microscope objective (1.15 NA, 40x). After passing through the imaging path of an inverted microscope the light beam is guided through an interferometer in common-path geometry and the off-axis interferogram is recorded by a CMOS-camera. (M1 − 4: mirrors, BS1 − 2: beam-splitters, L1 − 2: achromatic lenses, BEx: beam expander.

### 2.2. Reconstruction method

In our reconstruction algorithm we aim at reconstructing multiple-scattering samples, which undergo sustained rotation in the acoustic device, from a set of *L* fields 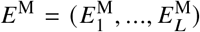 measured by off-axis interferometry. Since the rigid motion of the sample is not known *a priori*, our goal is to retrieve it together with the unknown object. Therefore, we need to solve the inverse problem

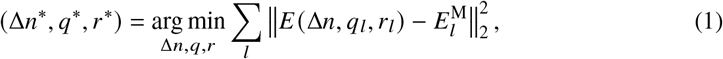

where *E* denotes the forward model, simulating the tomographic image formation for a refractive index contrast Δ*n*, rotation quaternion *q*_*l*_ and translation *r*_*l*_. Solving Eq. 1 blindly is highly challenging, as is well known from single particle cryo-electron microscopy [48]. Although the rotational trajectory of samples in our experimental setting is challenging to control precisely, it is not arbitrary; it possesses specific characteristics that we exploit: The sustained rotation induced by the acoustics has a clearly defined direction, i.e. the angular velocity does not change its sign. Additionally, the rotation tends to be smooth, and we assume no sudden angular changes in-between adjacent frames. We will capture the properties of the rotational trajectory with the regularization term 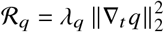, where ∇_*t*_ is the gradient operator with respect to time and *λ*_*q*_ a scalar parameter to tune the relative strength of the regularization. Additionally, we will make use of total variation regularization [12, 49, 50] for the object itself with 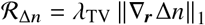, which has proven to be effective when solving inverse tomographic imaging problems. In total, we arrive at the following optimization problem to solve

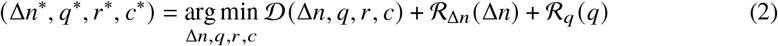

where the parameter *c* denotes additional unknowns in the experiments such as aberrations and tilts of the illuminating beam or the imaging plane and 𝒟 denotes the data-fidelity term (Eq. 1). To model the light-matter interaction including multiple scattering effects, we use multi-layer methods, which have been shown to accurately capture the transmitted light [10,12,14], even deep into tissues [51]. To solve Eq. 2 we rely on stochastic proximal gradient descent, where a batch of *b* images is processed for the calculation of every update of the variables. For initialization of the rotation, we utilize the approximate periodicity of the sample motion. Every rotation period is identified by cross-correlating the phase images. The rotation is initialized by a constant angular velocity during every rotation period and by a constant rotation axis determined by the acoustic actuation settings. For datasets consisting of multiple trajectories (separate recordings of the same object), e.g. if trajectories needed to be split or if two different rotational axes were recorded, the reconstruction algorithm will consist of multiple stages. In a first stage, Eq. 2 is solved independently for every continuously recorded trajectory, where the calibration parameters *c* are not included. In a second step, the relative orientation of the individual RI estimates are determined by a brute-force search in SO_3_ and further aligned by gradient descent. The reconstruction over the whole dataset, consisting of multiple continuous trajectories and including common calibration parameters *c*, is then initialized by the retrieved rigid-motion parameters and the average of RI distribution estimates processed in the first stage. For further details on the forward model and initializations we refer to Supplement 1. We implement the reconstruction algorithm in Python with JAX [52] and we leverage GPU acceleration using a Nvidia RTX 4090. Reconstruction times are given for the HEK293T cell for a volume of 150 × 174 × 200 from two times 2000 field images with a batch-size of 15. Individual reconstructions took about 150 − 200 iterations to converge, where one iteration over all 2000 images took ∼ 10 s, while the joint reconstruction converged in 200 additional iterations over the dataset of 2 × 2000 images, where the runtime for an iteration was ∼ 20 s.

## 3. Results

To evaluate the accuracy and explore the capabilities of our proposed approach, we conduct experiments on a set of different samples. We begin by assessing the accuracy of the RI reconstruction using bead samples, which have well-characterized RI and shape. Next, we apply our method to single cells, which are weakly scattering and commonly analyzed using ODT. To demonstrate the capabilities regarding the scalability of our method, we image 3D clusters of cells. Since spheroid growth can be well controlled and their scattering properties depend on maturation and size, these clusters serve as an excellent model for our investigations. Details on the sample preparation can be found in Supplement 1.

### 3.1. Validation with micro-bead clusters

First we test our method experimentally on clusters containing silica micro-spheres of known size of 3 µm and refractive index (1.43 − 1.46), which are levitated and rotated, either around a single axis or around two orthogonal axes. The results are shown in Figs. 3 and 4 with a 7-bead and a 3-bead cluster, respectively. The 7-bead cluster in Fig. 3 was rotated around one axis, and the reconstruction was performed with a dataset consisting of ∼ 1300 images. The initial estimate for the rotation quaternions, with a constant rotation velocity and the rotation axis oriented along the *z*-axis, is shown in Fig. 3 (b) in red color. The blue lines in Fig. 3 (b) depict the retrieved rotation quaternions, which clearly show variations in the angular velocity in addition to a tilted rotation axis. As part of the rigid motion of the sample, also the translation is retrieved, which is shown in the bottom of (b), where the yellow, magenta and green lines show the translations along the x, y, and z-axis, respectively. The refractive index reconstruction is shown as a 3D rendering in Fig. 3 (a), whereas Fig. 3 (c) shows slices through the retrieved refractive index distribution. As can be seen in (a) and (c), the spherical shape of the bead-cluster is well retrieved, as well as the RI values.

**Fig. 3.**
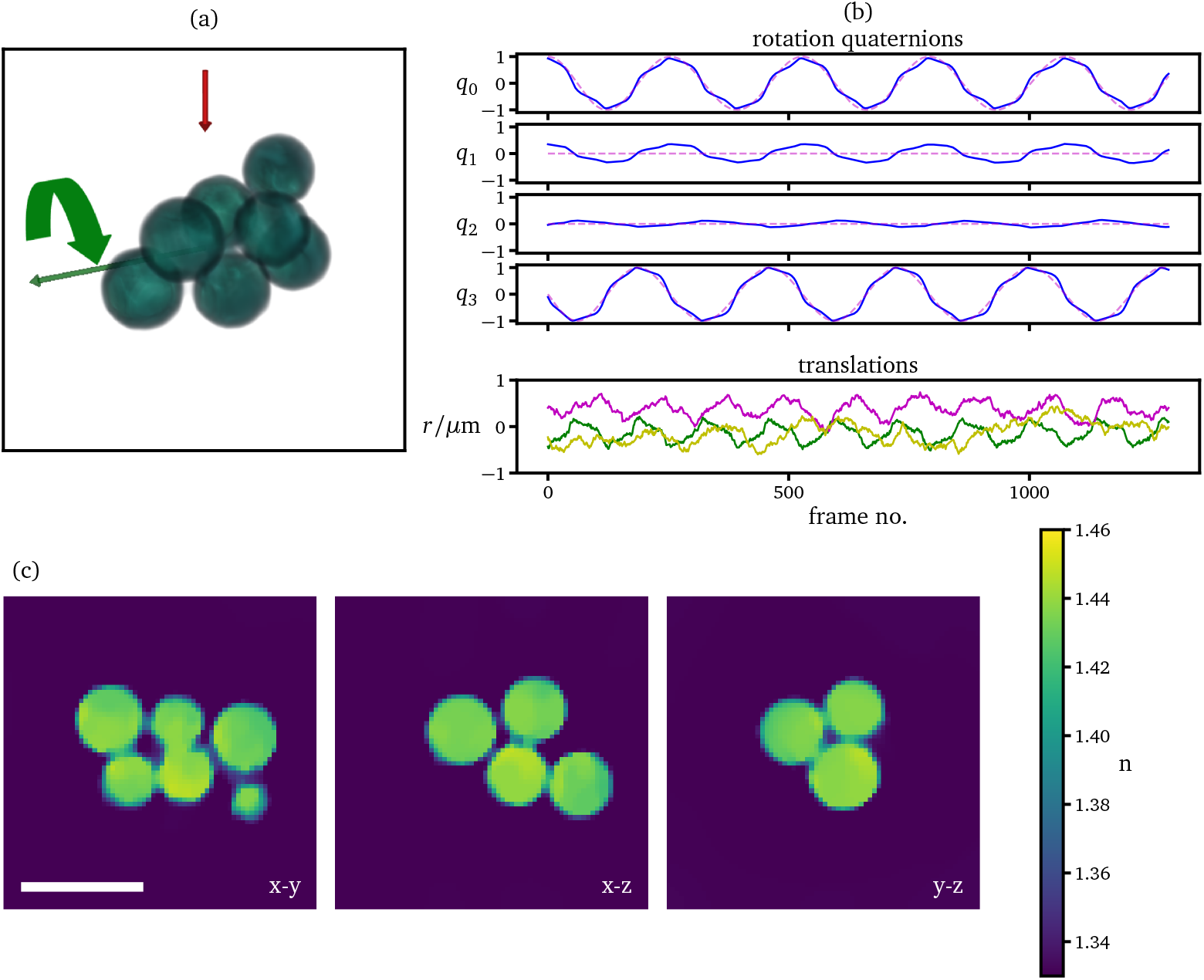
Reconstructions of a cluster of 3 µm silica beads rotated around a single axis: In (a) a 3D rendering of the retrieved refractive index distribution along with the rotation axis is shown. The red arrow depicts the illumination direction. In (b) the retrieved rotation quaternions are shown in blue, while the dashed magenta lines show the initial estimate with constant angular velocity and rotation axis. The images in (c) depict sections through the refractive index distribution. Scale bar: 5 µm.

**Fig. 4.**
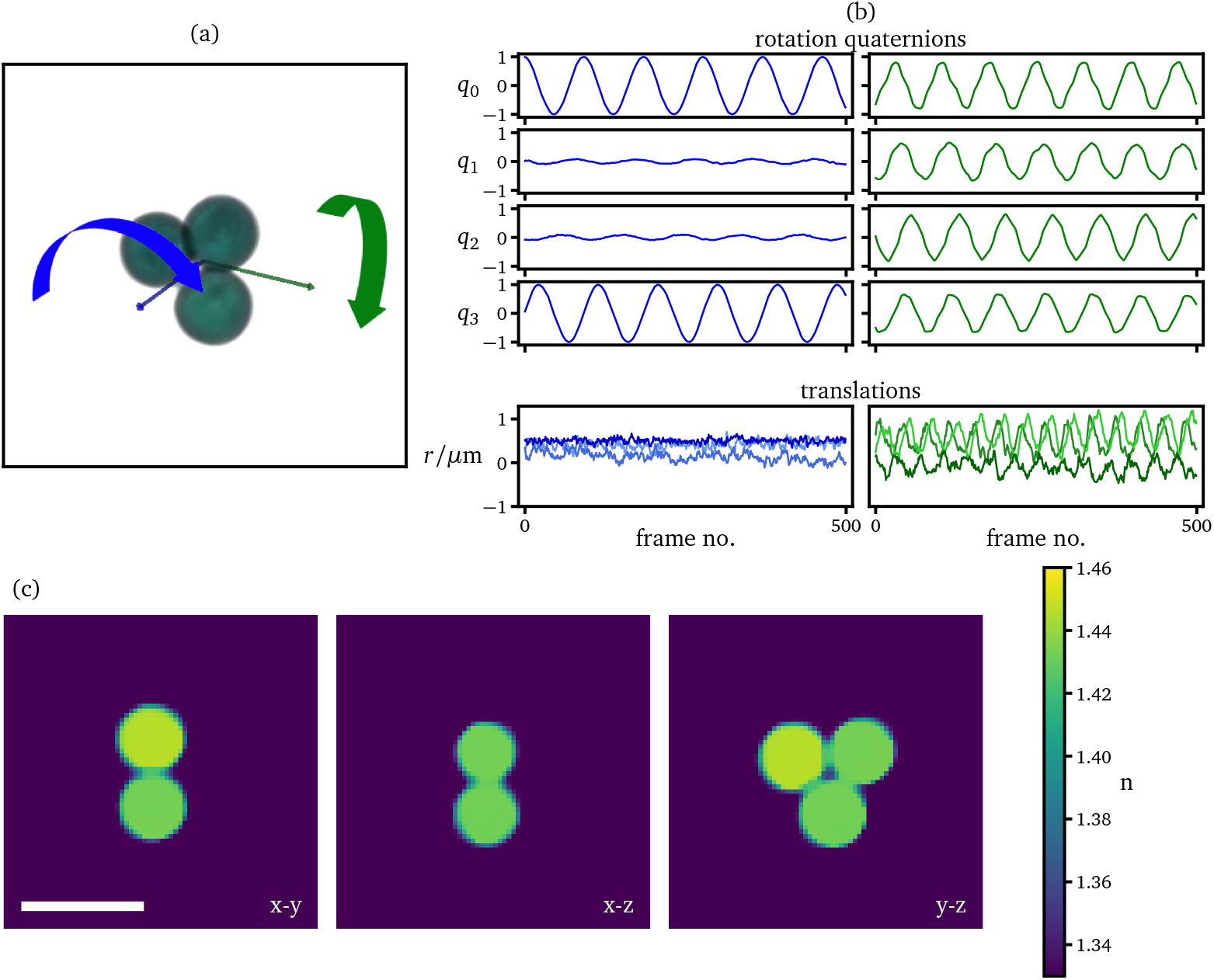
Reconstructions of a cluster of 3 µm silica beads rotated around two orthogonal axes: In (a) a 3D rendering of the retrieved refractive index distribution along with the rotation axes is shown. In (b) the retrieved rotation quaternions and translations are shown in colors that match the respective arrows in (a). The images in (c) depict sections through the refractive index distribution. Scale bar: 5 µm.

The trimer in Fig. 4 was rotated around *two* axes, which were oriented orthogonally with respect to each other, as well as with respect to the imaging axis. The reconstruction was performed first on the individual datasets, then we used the retrieved results for initialization of the common reconstruction using both datasets. The retrieved rotation quaternions of both trajectories are shown in Fig. 4 (b). The rotation around the minor axis of the object depicted in Fig. 4 (b) (blue arrow) shows a fairly uniform rotation compared to the rotation around the major object axis (green arrow). Due to the oblate-like shape of the three-bead cluster, the object is more (axis-)symmetric and has a more uniform mass distribution about the minor axis, hence the rotation is more regular. About the major axis, the object is less symmetric and hence the motion is more irregular as expected from different acoustic torque contributions acting on asymmetric samples in a not fully isotropic trapping potential [44, 45]. The three retrieved translation components are shown at the bottom of (b). In (a) and (c) a 3D rendering and slices through the retrieved refractive index distribution are shown. Compared to the retrieved RI in Fig. 3 with a single rotation axis, the retrieved RI using rotations around two orthogonal axes in Fig. 4 shows an increase in quality, where the retrieved RI distributions almost perfectly match the uniformity and spherical shapes, along with the sizes and RI values.

### 3.2. Biological results

We now move on to reconstructing biological samples of various complexity. Fig. 5 shows the reconstruction results of a kidney cell (HEK293T), which has been rotated around two approximately orthogonal axes. In Fig. 5 (a) a 3D rendering of the cell reconstruction is shown with the rotation axes. In (b) the retrieved rotation quaternions of both trajectories are shown. Similarly to the motion of the trimer in Fig. 4, the motion retrieved in Fig. 5 also shows stronger variations of the angular velocity for rotations around a major object axis. In Fig. 5 (c) we see slices through the retrieved 3D refractive index distribution with clearly visible structures, such as the nucleus containing nucleoli of higher refractive index. These structures are good test samples for checking that the resolution is truly isotropic. The availability of two orthogonal rotation axes greatly improves the reconstruction quality. To demonstrate this quantitatively, a comparison of the refractive index distributions retrieved with only a single rotation axis is included in Supplement 1. Since several methods perform reconstructions based solely on intensity images [9, 14, 19], we have also assessed what happens when the phase information is discarded, which required only minimal changes to the algorithm. A demonstration of reconstructions which rely on amplitude-only data along with a discussion regarding limitations in sample-size and imaging plane are provided in Supplement 1.

**Fig. 5.**
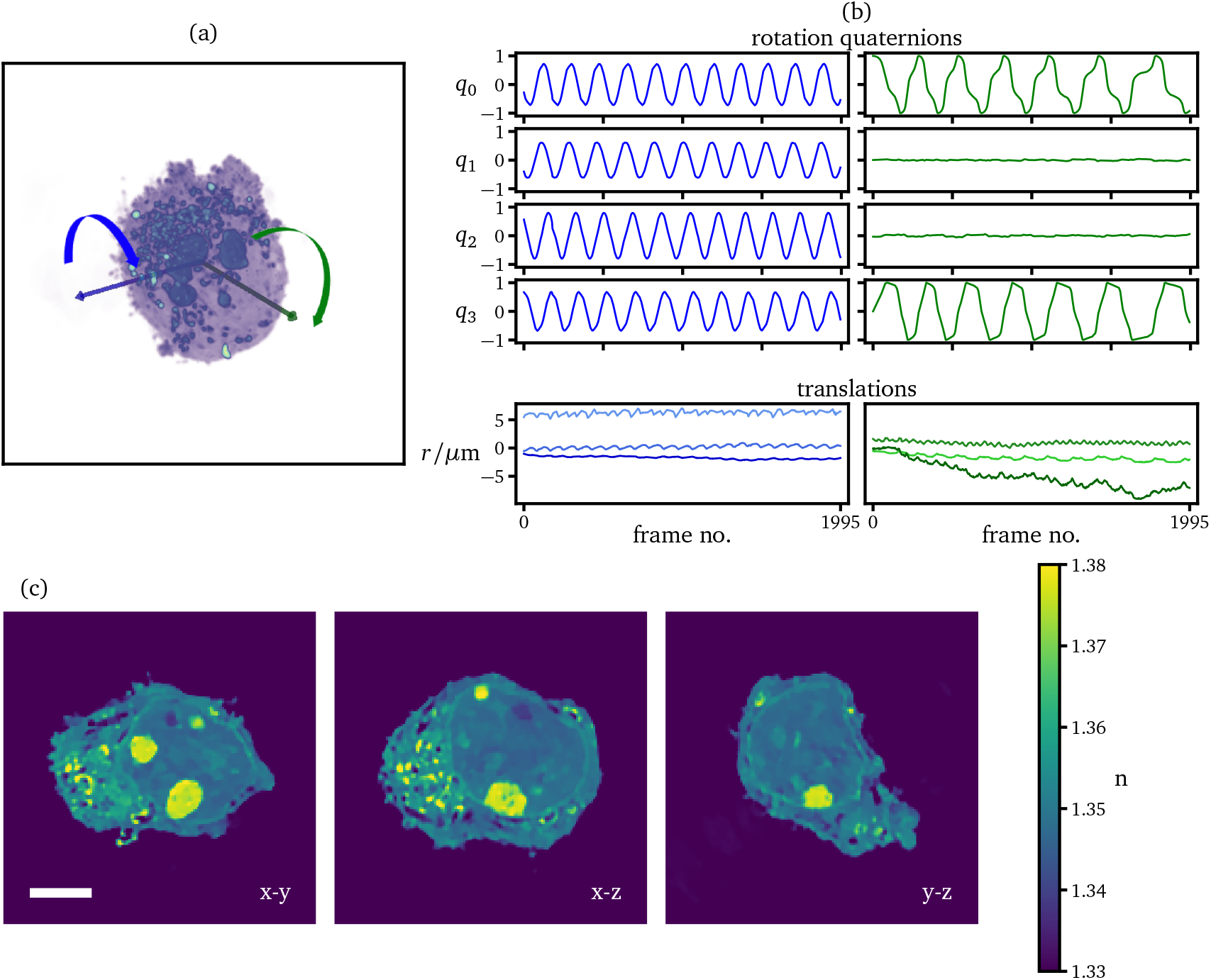
Reconstruction of an unstained HEK293T cell rotated around two axes: In (a) a 3D rendering of the retrieved refractive index distribution along with the rotation axes is shown. In (b) the retrieved rotation quaternions and translations are shown in colors that match those of the arrows in (a). The images in (c) depict sections through the refractive index distribution. Note that the outline of the nucleus with some nucleoli inside, is clearly visible. Scale bar: 5 µm.

We now tackle the most challenging type of samples, optically dense cancer cell clusters containing about 100-150 cells estimated by volume. One difficulty when performing ODT on such large specimens is the presence of local minima during optimization. To avoid this, it has proven useful to initialize the individual reconstructions first with a projection-based forward model on the phase part of the measurement, which has been unwrapped [53]. For details on the initialization we refer to Supplementary section 2. Figures 6 and 7 show results of unstained Neuroblastoma cancer cell clusters. In Figs. 6 and 7, (a) depicts a 3D rendering with the rotation axis, whereas (d) shows sections through the refractive index distribution. In (b) the retrieved rotational and translational trajectories are shown. The cancer cell clusters in both Fig. 6 and 7 were rotated repeatedly around approximately the same rotation axis. We see that the retrieved rotational trajectories exhibit very similar shapes. In (c) we see additionally the retrieved Zernike coefficients along with the aberration map in the pupil, where the first 5 radially symmetric Zernike polynomials were chosen as basis. We found that the presence of small and strong scatterers enable the retrieval of aberrations due to the scattering at high angles. In Figs. 6 and 7 (d) we see the individual cells as well as the subcellular structure clearly resolved. Additionally, high refractive index structures correlate with lipid droplets and are sharply imaged throughout the entire volume, which is a strong indication that the algorithm indeed converged to a valid rotational trajectory close to the ground truth. Fluorescence analyses were performed to verify lipid droplet structure and localization as well as larger nuclear areas, which are provided in Supplement 1. Three dimensional renderings as well as flythroughs through the retrieved RI distributions of the cell clusters depicted in Figs. 6 and 7 are provided in Visualizations 1-4. We are not aware of any other ODT reconstruction method that can currently provide this kind of quality for such a sample, at least as far as isotropic resolution is concerned.

**Fig. 6.**
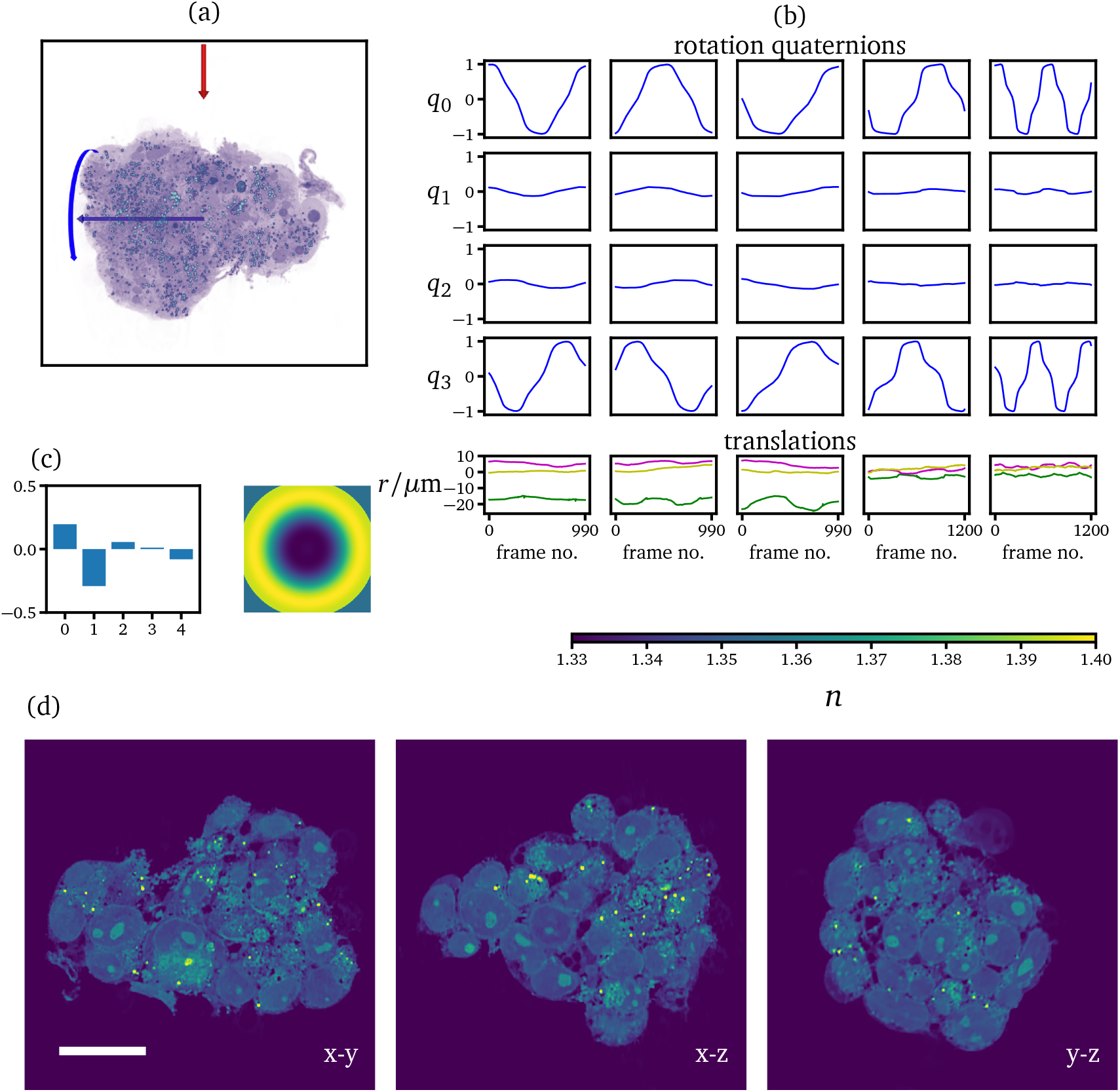
Reconstruction of an unstained Neuroblastoma cell cluster: In (a) a 3D rendering of the retrieved refractive index distribution along with the rotation axis is shown. In (b) the retrieved rotation quaternions are shown in blue. In (c) the retrieved Zernike coefficients (first 5 radial Zernike) along with the aberration map is shown. The images in (d) depict sections through the refractive index distribution. Scale bar: 20 µm.

**Fig. 7.**
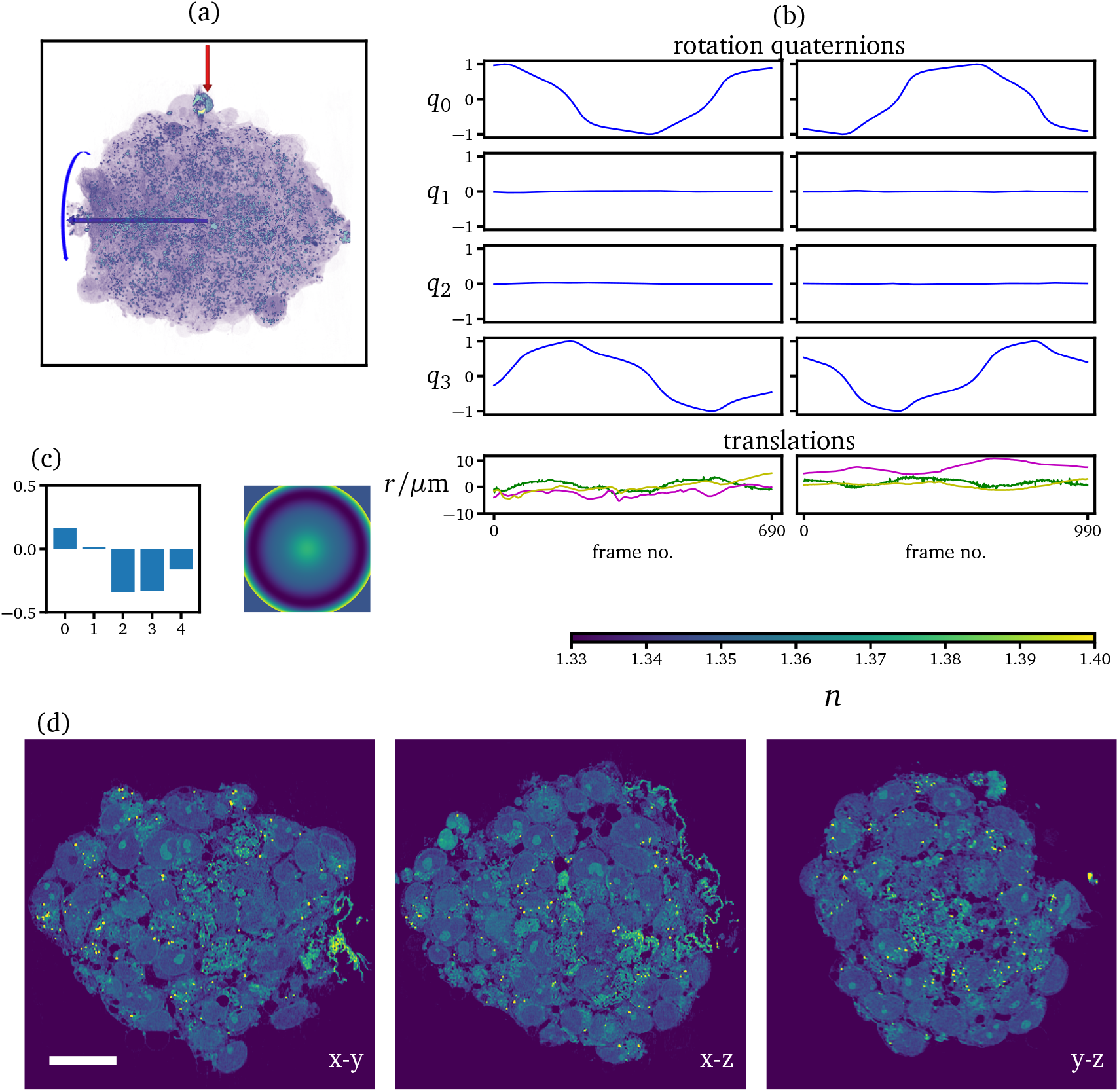
Reconstruction of an unstained Neuroblastoma cell cluster rotated along two trajectories: In (a) a 3D rendering of the retrieved refractive index distribution along with the rotation axes is shown. In (b) the retrieved rotation quaternions are shown in blue, along with the retrieved translations. In (c) the retrieved Zernike coefficients (first 5 radial Zernike) along with the aberration map is shown. The images in (d) depict sections through the refractive index distribution. Scale bar: 20 µm.

## 4. Discussion

A fundamental challenge in optical microscopy is the missing cone problem, which arises from inaccessible spatial frequency components due to the limited NA of illumination and detection optics. While rotating the object gives access to these missing frequencies, the critical issue lies in accurately controlling or retrieving the rotational motion. In this work, we presented an approach to recover both object and motion in a consistent manner, which relies on gradient-based optimization. An acoustic strategy provided contact-less and non-invasive rotational actuation of the object, while a common-path interferometer enabled the recording of phase and amplitude.

### Limitations

A potential limitation of the presented approach lies in its reliance on nonlinear optimization, where convergence to the global optimum is not guaranteed. This challenge is also shared by other methods, which incorporate multiple-scattering and introduces additional complexity when recovering both object and motion. Clearly, the computational cost of a joint object and motion retrieval is higher compared to algorithms that focus solely on object recovery. Furthermore, for objects with symmetries or without inner structures, retrieving object and rotations simultaneously can be become severely ill-posed. To mitigate these issues, we employed suitable initialization and regularization methods for both object and motion. Future work could involve the formulation of physics-based priors on the rotational and translational trajectories. Furthermore, applying deep-learning based approaches to the recovery of the rigid-motion such as in [54] could provide better initial estimates for the rotational trajectory.

### Forward model trade-offs

Modeling the light matter interaction is subject to the trade-off between accuracy and computation time. Coherent propagation methods which rely on the single-scattering approximation are fast, but fail to accurately capture the scattering process for larger samples and limit the applicability of conventional ODT. Thus, for challenging samples even the next higher order approximation beyond 1st order Born or Rytov, such as the common-circles method [55], will not provide a reliable reconstruction. On the other hand, solving the full Helmholtz equation in a volumetric fashion has shown to increase the fidelity of reconstructions, particularly in the illumination angle-scanning configuration [18, 56, 57]. However, using these models comes at a greater computational burden, and the full capability of these models is not utilized in ODT, since the reflected light components are typically not recorded. Regarding accuracy and computation time, multi-layer propagation models occupy a spot in-between single-scattering and full solutions to the Helmholtz equation. They represent a key tool for the success of the presented approach due to their ability to model multiple-scattering in the forward direction, while keeping a low computation time. As the light propagation remains orthogonal to the imaging plane, the problem of obliquity [15, 19, 58] does not introduce errors. Furthermore, the quality of the reconstructions demonstrates that these multi-layer methods are able to accurately model the multiple-scattering process also deep into tissue, which limits conventional ODT. Recently, efficient methods for imaging deep into specimens for ODT have been developed [59, 60], which could also represent viable candidates for a forward model.

### View diversity

An important factor regarding the tractability of the inverse problem, as well as the overall quality of the reconstruction, is given by the view diversity which is contained within the rotational trajectories. Multiple trajectories, which rotate approximately around the same axis, do improve the tractability of the problem, but also contain redundancy. In contrast, using multiple rotational trajectories which are oriented orthogonally with respect to each other contain a greater view diversity. Thus, methods capable of performing rotations around multiple orthogonal axes – such as our proposed approach based on acoustic actuation [44, 45], along with dielectrophoresis [37] and fiber-optical traps [61] – are especially well suited and desired for the proposed reconstruction method.

Compared to mechanical mounting of objects, label-free and non-contact rotation techniques can be less invasive, simplify sample preparation and render the sample more accessible - not only with regards to accessible imaging angles, but also for potential downstream manipulation or analysis, as for instance in microfluidic devices for sorting. The presented approach can also be readily applied in situations where the samples are rotated by flow-based approaches, optical methods, e.g. fibre-optical traps or optical tweezers, or further acoustic strategies, as long as the motion is continuous. Furthermore, reconstructing 3D object and motion of a sample rotated by acoustic actuation enables estimating the forces and torques exerted on the particle by the acoustic pressure field as well as the viscous forces of the fluid. In future studies the joint retrieval of object and motion could therefore be used to infer mechanical and elastic properties of cells and tissues. As generating view-diversity through object rotation provides a solution to anisotropic spatial resolution, which inherently plague imaging methods, rotational manipulation strategies along with reconstruction approaches are much sought-after. As for imaging methods a formulation of physical forward models are available, the presented approach can be applied to a variety of imaging modalities. Furthermore, as refractive index variations induce refraction, diffraction and scattering in biological specimens, they are highly relevant also for fluorescent techniques. Knowledge of the refractive index distribution enables the digital correction of scattering, which opens possibilities for fluorescent imaging deep inside specimens [62–64].

## 5. Conclusion

In this work we presented an approach for optical tomographic imaging of multiple-scattering samples with rotational trajectories that are not known precisely *a priori*. We introduced a computational model that is able to retrieve both, the object and its position and orientation, from phase-sensitive measurements recorded while the object is undergoing sustained rotation by acoustic force-field actuation in a non-contact way. We performed high-resolution reconstructions of clusters of silica microspheres of known properties, of individual cells and of dense cancer cell clusters containing more than 100 cells.

The method presented here extends optical tomography into a regime of biological samples that was hitherto inaccessible to ODT. Besides being capable of providing fully *isotropic* resolution, as a by-product also the time-dependent motion of the sample in 3D is retrieved, for further analysis. The approach is not limited to acoustic levitation and actuation, but is applicable to a variety of situations where motion-uncertainty is inevitable. We herewith introduce an enabling tool for studying developing *in vitro* models in a biocompatible, low-impact manner, which has the potential to benefit various fields in biomedical and material science research.

## Supporting information

Visualization 1

Visualization 2

Visualization 3

Visualization 4

## Acknowledgment

S.M. would like to thank Prof. Demetri Psaltis for fruitful discussions. We thank Armin Sailer (Institute for Experimental Physics, University of Innsbruck) for CNC-manufacturing the high-NA chip according to our design. We thank Prof. Alexander Jesacher for helpful discussions and suggestions.

## Data availability

Data underlying the results presented in this paper are not publicly available at this time but may be obtained from the authors upon reasonable request.

## Supplemental document

See Supplement 1 for supporting content.

