## Supplement 1 for "Optical tomography reconstructing 3D motion and structure of multiple-scattering samples under rotational actuation"

### 1. FORWARD MODEL

#### A. Multi-layer model

To propagate a wave-field through a 3D refractive index contrast distribution $\Delta n \in \mathbb{R}^{n_x \times n_y \times n_z}$ given on a cartesian grid with size $(n_x, n_y, n_z)$ we subdivide $\Delta n$ along the optical axis (here z-axis) into layers and propagate an initial field $E_0 \in \mathbb{C}^{n_x \times n_y}$ through these sequentially. This approach has proven to be very effective to model the tomographic image formation for multiple-scattering samples in transmission. For on-axis propagation, the models [1–4] have similar accuracy. Thus, we chose the Beam Propagation Method or multi-slice method [2], since it is the simplest and most efficient multi-layer method. It consists of alternating between a free-space propagation $K(E) = \mathcal{F}^{-1}[\mathcal{F}[E]\exp(ik_z\Delta z)]$ and a phase modulation step $P(E) = E\exp(ik_0\Delta n\Delta z)$, where $\mathcal{F}$ and $\mathcal{F}^{-1}$ denote the forward and inverse 2D Fourier transforms. The quantity $k_z$ denotes the $z$-spatial frequency component component and is given by $k_z = \sqrt{(k_0 n_0)^2 - k_x^2 - k_y^2}$ with $k_x$ and $k_y$ denoting the spatial frequency components in $x$ and $y$ direction, and $k_0 = 2\pi/\lambda_0$ where $\lambda_0$ is the wavelength in vacuum and $n_0$ the refractive index of the background medium. To compute the refractive index distribution at rotated and translated coordinates we utilize linear interpolation.

#### B. Aberrations

We assume our imaging system to be linear and shift invariant, which allows us to model aberrations as a single kernel in spatial frequency space. To model the aberrations we chose the Zernike basis $Z_m^n$, where we found it sufficient to consider only even orders in $n$ with $m = 0$, resulting in radially symmetric aberrations $W = \sum_i c_i Z_0^{2i+2}$. In our forward model, we consider the position of the imaging plane as a separate parameter, such that we subtract the contribution $k_z = \sqrt{(k_0 n_0)^2 - k_x^2 - k_y^2}$ from the calculated aberration

$$W_{\text{ref}} = W - \frac{< k_{z,n}, W_n >}{< k_{z,n}, k_{z,n} >} k_z \tag{S1}$$

where the vectors with subscript $n$ define the mean-subtracted vector $v_n = v - 1/N \sum_i v_i$ and the brackets $< \cdot, \cdot >$ denote the inner product.

### 2. RECONSTRUCTION

The objective function that we minimize is of the form

$$\min_{\Delta n, q, r, c} \mathcal{D}(\Delta n, q, r, c) + \mathcal{R}_{\Delta n}(\Delta n) + \mathcal{R}_q(q) \tag{S2}$$

using gradient based optimization with a "mini-batch" stochastic optimization strategy. Our approach for updating the optimization variables involves selecting indices $i_k$ randomly from the dataset (without replacement) and updating $\Delta n$ and $c$, while accumulating the gradient parts of rotation quaternions $q$ and the translations $r$

$$\Delta n_{k+1} = \text{prox}_{\alpha_{\Delta n}\mathcal{R}_{\Delta n}}(\Delta n_k - \alpha_{\Delta n}\nabla_{\Delta n}\mathcal{D}_{i_k}) \tag{S3}$$

$$c_{k+1} = c_k - \alpha_c \nabla_c \mathcal{D}_{i_k} \tag{S4}$$

$$(\bar{q}_{n+1})_{i_k} = \nabla_q \mathcal{D}_{i_k} \tag{S5}$$

$$(\bar{r}_{n+1})_{i_k} = \nabla_r \mathcal{D}_{i_k}, \tag{S6}$$

where $\mathcal{D}_{i_k} = \sum_{i \in i_k}$. After a full pass through the dataset, we update the rigid motion parameters

$$q_{n+1} = \text{prox}_{\alpha_q \mathcal{R}_q}(q_n - \alpha_q \bar{q}_n) \tag{S7}$$

$$r_{n+1} = r_n - \alpha_r \bar{r}_n \tag{S8}$$

where $n$ denotes the number of total passes through the dataset. We therefore apply a stochastic iteration scheme for the parameters $\Delta n$ and $c$, while the gradients for $q$ and $r$ are updated once for every pass through the dataset. To evaluate the proximal operator $\text{prox}_{\alpha_{\Delta n}\mathcal{R}_{\Delta n}}$ we utilize [5, 6], while $\text{prox}_{\alpha_q \mathcal{R}_q}$ has a closed-form solution.

In the reconstruction of samples from multiple data-sets or 'time-series' (e.g. separate recordings) we follow two stages. In the first stage, the reconstructions are first performed independently for each time-series, yielding two estimates of the refractive index distribution. The relative orientation between the two estimates is determined by a brute-force search of the whole $SO_3$ by random search, for which we utilize [7] to generate the quaternions.

The random rotation which yielded the highest cross-correlation was then further refined by a gradient descent, where we solve

$$\arg\min_{q,r} \|\Delta n_1 - \Delta n_2(q,r)\|_2^2 . \tag{S9}$$

In the brute-force search we chose 1000 random rotations, which gave an estimate that was sufficiently close as initialization for the refinement in all datasets. The final reconstruction is then initialized by the average of all the individual reconstructions and considers a multiple time-series. The continuity of the rotation trajectory is then enforced for each time-series independently, while the aberration is assumed to be equal among all time-series.

Fig. S1 shows the individual reconstructions of the HEK-293T cell in (a) and (b), which have undergone rotations around approximately orthogonal axes. Both reconstructions in (a) and (b) exhibit artifacts and blurring. However, the shape is registered well enough to retrieve the relative orientation, which is then used to initialize the object and motion for common reconstruction shown in (c).

### 3. INITIALIZATION

The recorded off-axis holograms undergo standard pre-processing by isolation of the first order in Fourier space. For every time-series we additionally record a hologram without the sample and divide the reconstructed fields by this reference field to get rid of residual phase-errors in the reconstruction.

To obtain an initial estimate of the rotation angles, we calculate the cross-correlation between the first image and the time-series. The peaks of time series identify the rotational period. For initialization we assume a constant angular velocity during each rotation period. The rotation axis is initialized as constant over the whole time-series and is chosen either in the $x$ or $y$ direction depending on the acoustic actuation parameters.

#### A. Cancer spheroids:

For the cancer spheroids we performed an additional step to initialize the wave-optical reconstruction. Starting from an uninitialized object, i.e. $\Delta n = 0$ is appropriate for single cells, but tends to fail in cases where the accumulated phase exceeds multiples of $2\pi$, due to local minima. In these cases we performed phase-unwrapping on the measured phase images [8]. An estimate of the refractive index along with rotation and translation is then calculated by solving a reduced optimization problem

$$\arg\min_{\Delta n,q,r} \sum_l \left\| R(\Delta n, q_l, r_l) - P_{\text{unwrapped},l} \right\|_2^2 + \lambda_{\text{TV}} \|\Delta n\|_{\text{TV}} + \lambda_q \|\nabla_t q\|_2^2 , \tag{S10}$$

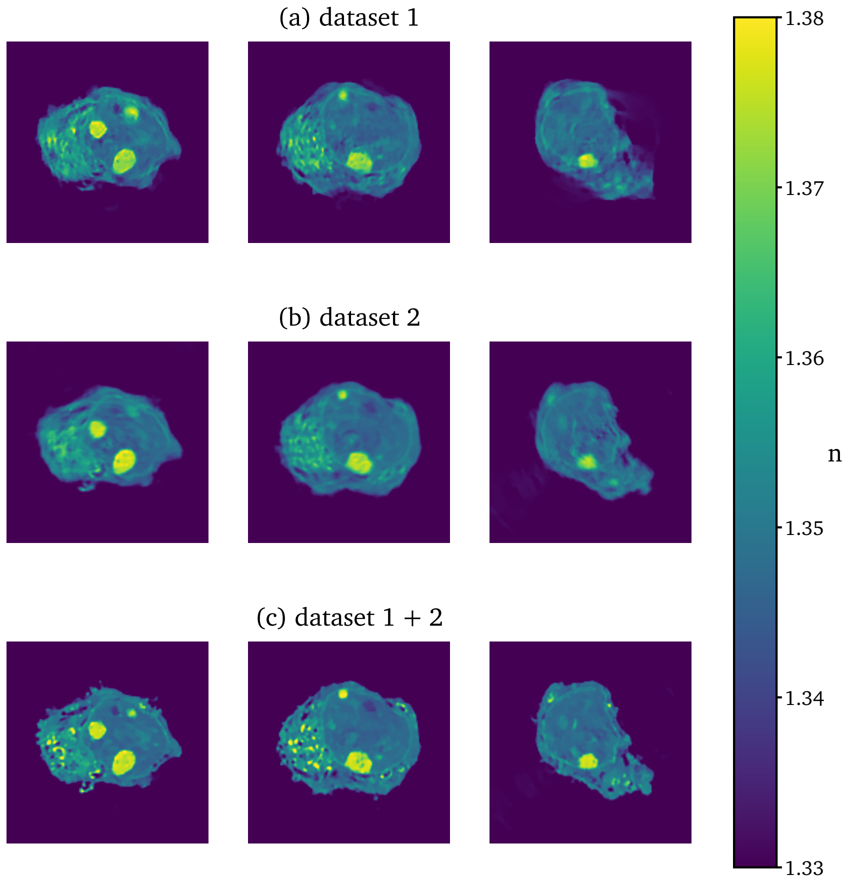


**Fig. S1.** Individual and reconstructions: (a) and (b) show the reconstructions of a HEK-293T cancer cell where only rotations around a single axis were considered, whereas (c) shows the joint reconstruction performed in two stages.

where the forward model $R(\Delta n, q_l, t_l)$ denotes the Radon-transform and $P_{\text{unwrapped},l}$ the unwrapped phase images. Eq. S10 is then solved in a similar fashion as Eq. S2. Although the unwrapping process tends to show errors at multiple positions in each image, the reconstruction from the whole time-series was of sufficiently high quality for initialization with the wave-optical model.

### 4. INTENSITY ONLY INFORMATION

We now discuss the outcome if one relies solely on the amplitude of the recorded fields for reconstruction. This simplifies the experimental setup, as phase-stable measurements are no longer necessary. The reconstruction algorithm can also be adapted to intensity only information by simply exchanging the error metric to

$$(\Delta n^*, q^*, r^*, c^*) = \underset{\Delta n, q, r, c}{\arg\min} \sum_l \left\| |E(\Delta n, q_l, r_l, c)| - \sqrt{I_l^{\mathrm{M}}} \right\|_2^2 + \mathcal{R}_{\Delta n}(\Delta n) + \mathcal{R}_q(q) \tag{S11}$$

where $I_l^{\mathrm{M}}$ denotes the measured intensities. However, in the absence of phase measurements the focal plane at which the images are recorded has an influence on the refractive index distribution. Reconstructions using intensity only information are shown in Fig. S2, where (a) depicts the reconstructions for intensity images, where the focal plane is located approximately at the rotational center of the cell, whereas the reconstruction in (b) utilizes intensity images recorded $\sim 15$ µm away from the rotational center. For reference, Fig. S2 (c) depicts the reconstructions with amplitude and phase. We can recognize that the reconstruction results differ considerably from each other, where the reconstruction from amplitude images located at a distance from the center (b) is closer to (c). Additionally, the motion recovered in (b) is very close to the motion recovered in (c), whereas recovery of rotational parameters in (a) was unstable. Furthermore, we see that the reconstructions with amplitude-only information have a more pronounced background, which is absent in the reconstruction from amplitude and phase. We therefore conclude that the method also works when only the amplitude information is available, although some care has to be taken in the choice of the focal plane, where the images are recorded.

### 5. SAMPLE PREPARATION

**Bead clusters.** Clusters of beads were created by mixing a monodisperse suspension of 2.9 µm ($\pm 0.4$) µm silica beads (Whitehouse Scientific $^{\text{TM}}$) with isopropanol. This suspension was distributed onto microscope slides, and after allowing the isopropanol to evaporate completely, the resulting dry mass was scraped off and subsequently redistributed in water. The process ensured that the beads formed rigid bonds with one another within clusters of various sizes.

**Cell lines.** HEK 293T (CRL-3216) cells were purchased from ATCC and cultured in EMEM (Merck, Vienna, Austria) and DMEM (Merck, Vienna, Austria), respectively. The neuroblastoma line STA-NB15 [9, 10] were cultured in RPMI1640 (Merck, Vienna, Austria). All media contain 10% fetal bovine serum (Thermo Fisher Scientific, Waltham, USA), 100 U/ml penicillin, 100 µg/ml streptomycin, and 2 mM L-glutamine (Merck, Vienna, Austria). All cultures were routinely tested for mycoplasma contamination using the Universal Mycoplasma Detection Kit (ATCC LGC Standards GmbH, Wesel, Germany).

**Hanging drops.** NB15 cells were counted and plated at 15-20 cells per 40 µl drop on the lid of conventional cell culture dishes. Evaporation was prevented by PBS in the lower chamber. At a size from 40 – 80 µm (approximately after 48 to 60 hours), spheroids were transferred into V-Bottom plates (Thermo Fisher Scientific, Waltham, USA), and fixed with 4% HistoFix solution (Carl Roth, Karlsruhe, Germany) for further experiments.

**Live cell fluorescence microscopy.** Cells were grown on glass slides or LabTek Chamber SlidesTM (Nalge Nunc International, USA) coated with 0.1 mg/ml collagen for live cell fluorescence analyses. Non-attached cells were imaged immediately after seeding, adhered cells were imaged 24 hours after seeding. Lipid droplets were stained with 1µg/ml Bodipy 493/503 (Thermo Fisher Scientific, Waltham, USA) in PBS according to manufacturer instructions, nucleoli were stained with a Nucleolar-ID Green detection Kit (Enzo Life Sciences, Lausen, Switzerland) according to manufacturer instructions. Nuclei were visualized with 500ng/ml Hoechst33342 (Merck, Vienna,

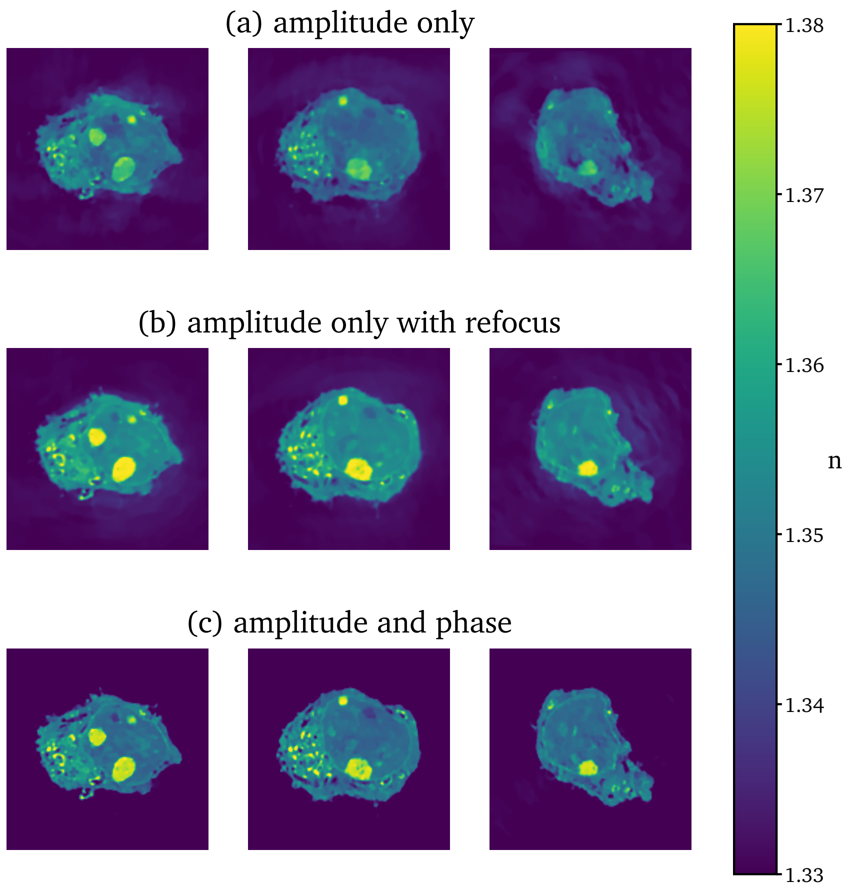


**Fig. S2.** Comparison with amplitude-only reconstruction of a HEK293T cell: (a) depicts the reconstruction from amplitude only images, where the focal plane is located around the rotational center, whereas (b) shows reconstructions from amplitude only images with the focal plane at a $\sim$ 15 μm from the rotational midpoint. In (c) the reconstruction from full field data is depicted.

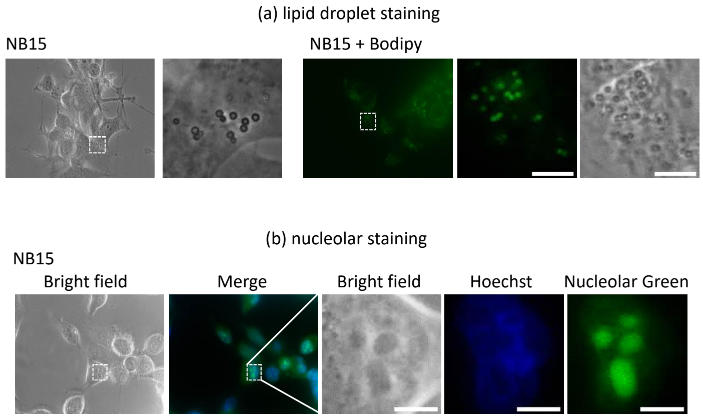


**Fig. S3.** Staining of lipid droplets and nucleoli: (a) NB15 cells were stained with 1μg/ml Bodipy 493/503 in PBS to visualize lipid droplets (green). (b) NB15 cells were incubated with NUCLEOLAR-ID® Green detection kit in assay buffer according to manufacturer's instructions. DNA (blue) was counterstained using the DNA-intercalating dye Hoechst33342 (500ng/ml). Scale bar is 5 μm.

Austria). All images were acquired using an Axiovert200M microscope and analyzed in the Axiovision Software (Zeiss, Vienna, Austria).

Small dots correlate with the size and localization of lipid droplets. Fluorescence analyses using the green fluorescent dye Bodipy 493/503 were performed to verify lipid droplet structure and localization as shown in Fig. S3 (a). Larger, nuclear areas are most likely nucleoli as suggested by staining cells with Nucleolar-ID Green detection kit and fluorescence microscopy, Fig. S3 (b).

### 6. ACOUSTIC TRAPPING

The geometry, characterization and operation of the acoustofluidic device is described in detail in [11]. In short, the three orthogonal main channels – two long crossed horizontal channels and one short vertical channel – and the microfluidic channels are CNC machined in an aluminium plate. A transducer is attached to each channel as seen in Fig. 2 in the main article. The optically transparent transducers are made of lithium niobate ($LiNbO_3$) (36° Y-cut, 1 mm thickness, Roditi) with transparent ITO electrodes (sheet resistance 10Ω/sq, Diamond Coatings), with a thickness resonant mode frequency of the bare transducer around 3.4 MHz and a wavelength in water of about 440 μm. A coverslip seals the bottom of the chamber and also acts as the reflector of the waves from the vertical transducer, while steel reflectors are inserted into the end of the horizontal channels. The transducers are operated in thickness mode and the bulk waves generate standing waves upon reflection. We optimize the chip geometry to support resonant acoustic modes in the vertical and horizontal directions at the same frequency.

The chamber is filled with water and placed on the microscope stage. With the transducers active, the sample suspension is introduced via the microfluidic channel leading the sample below the vertical transducer to the region where all three acoustic waves intersect and where we have optical transmission access. The sample is levitated and trapped in a pressure node due to the standing wave generated by the vertical transducer, and with the additional waves from the horizontal transducers we further confine the sample in 3D and have individual control of the acoustic radiation forces in each direction.

We control the rotational manipulation by adjusting the magnitude between the acoustic radiation torque contributions in our device. To transiently rotate a sample, we change the relative amplitude between the standing waves, and the acoustic scattering by an extended asymmetric object results in an acoustic restoring torque that aligns the particle to the new acoustic force landscape. To induce a sustained rotation, we generate specific acoustic modes which can continuously transfer angular momentum to an object via viscous dissipation in the fluid and object [12]. We generate the spinning torque by exciting two orthogonal standing waves at the exact same frequency at a non-zero phase shift [13, 14]. In order to induce sustained rotations around an axis orthogonal to the optical axis, we generate the spinning torque by operating the vertical transducer and one horizontal transducer (propagating along y-direction, as an example). A sustained rotation will occur when the spinning torque is larger than the restoring torque, which we achieve by tuning the relative phase between these two transducers and also minimizing the restoring torque by adjusting their relative amplitude to create a largely isotropic trapping potential (in the YZ-plane in this case). An object will then rotate around an axis orthogonal to the two standing waves generating the spinning torque (around x-axis in this case). The third transducer (with propagation along x-axis) is operated at a different frequency not to influence the spinning torque, and is kept at a low voltage relative to the two other transducers. The object will align its major axis parallel to the direction of weakest trap stiffness (along x-axis), hence it will rotate around its major axis.

Now, to induce a sustained rotation around the orthogonal object axis, we increase the amplitude of the third transducer so the trap stiffness is the largest in this direction (along x-axis). In response to this restoring torque, the object will realign with its minor axis parallel to the x-axis. However, the the spinning torque is unchanged and still leads to a rotation around the the x-axis, hence the object will now rotate around its minor axis.

We choose a resonance frequency close to the 1st, 3rd or 5th harmonic frequency of the transducers to operate close to the regime where the targeted sample diameter, d $\lessapprox \lambda/3$ for sufficient trap stiffness. For single cells (around 10 µm to 30 µm in size), we operate at the 5th harmonic frequency ($\approx$ 20 MHz), and for the used spheroids (around 40 µm to 100 µm in size) at the 1st or 3rd harmonic frequency ($\approx$ 3 MHz or 11 MHz, respectively). The transducers are driven by a sinusoidal signal by waveform generators: one dual output signal generator (Keysight 33522B) and one single output generator (Agilent 33220A). The transducers are operated in the range of 3 V to 20 V peak to peak.
